# Dynamic Palmitoylation Events Following T-Cell Receptor Signaling

**DOI:** 10.1101/831388

**Authors:** Eliot Morrison, Tatjana Wegner, Andres Ernesto Zucchetti, Miguel Álvaro-Benito, Ashley Zheng, Stefanie Kliche, Eberhard Krause, Britta Brügger, Claire Hivroz, Christian Freund

## Abstract

Palmitoylation is the reversible addition of palmitate to cysteine via a thioester linkage. Following stimulation of the T-cell receptor we find a number of proteins are newly palmitoylated, including those involved in vesicle-mediated transport and Ras signal transduction. Among these stimulation-dependent palmitoylation targets are the v-SNARE VAMP7, important for docking of vesicular LAT during TCR signaling, and the largely undescribed palmitoyl acyltransferase DHHC18 that is expressed in two isoforms in T cells. Using our newly developed On-Plate Palmitoylation Assay (OPPA), we show DHHC18 is capable of palmitoylating VAMP7 at Cys183. Cellular imaging shows that the palmitoylation-deficient protein fails to be retained at the Golgi.

---

The initial signaling events following the recognition of a peptide-MHC complex on an antigen-presenting cell (APC) by the T-cell receptor (TCR) involve a number of important molecules that must localize to the plasma membrane (PM) at the immunological synapse (IS). Palmitoylation, the reversible linking of 16-carbon palmitic acid to cysteines via a labile thioester bond, is one strategy used by several important TCR-signaling proteins to drive this localization to and from the membrane (e.g. the soluble Src-family kinases Lck and Fyn) or to and from liquid-ordered microdomains (“lipid rafts”) within the membrane (e.g. the transmembrane scaffold LAT)^1^.

Unlike other lipid modifications like myristoylation or prenylation, the reversible chemistry of the thioester bond allows palmitoylation to be both added to and removed from specific cysteines; indeed, enzymes have evolved to both add (e.g. the DHHC family of protein acyltransferases [PATs]) and remove (e.g. acyl protein thioesterase 1 [APT1] and ABHD17) palmitate from these sites in eukaryotes^2,3^. This has given rise to palmitoylation cycles, where proteins are palmitoylated in the ER/Golgi, cycled to the PM via vesicle trafficking and localized to the site of activity, then depalmitoylated and recycled back to the ER/Golgi^4^.

Such cycles have become well-established in neuronal proteins like PSD-95, as well as the G-protein α subunits Gα_i_ and Gα_q_ and small GTPases H-Ras and N-Ras^4,5^, and pulse-chase experiments have shown that palmitoylation turnover can be accelerated following signaling stimuli, such as glutamate receptor activation with PSD-95, GTP binding of H-Ras, and GPCR signaling for Gα proteins^6–8^. In the context of T-cells, Lck has been shown to undergo relatively rapid palmitate turnover under resting conditions, which is accelerated following Fas receptor stimulation, and this palmitoylation is specifically catalyzed by DHHC21 at the PM^9^. A decrease in the palmitoylation of LAT has been observed in anergic T cells, which affects its ability to localize to lipid rafts and induce an effective stimulatory response^1,10^. Docking of vesicular LAT to the IS and downstream signaling also critically depend on palmitoylated SNARE proteins like SNAP-23 and VAMP7^11,12^. While these targeted approaches suggest the possibility of regulated palmitoylation cycles, a global approach tracking changes of the complete palmitome of T-cells following a stimulatory event has not been reported.

Equally enigmatic are the enzymes responsible for catalyzing the palmitoylation reaction, the DHHC family of PATs, first identified in yeast more than 15 years ago^13,14^. While 23 PATs are expressed in mammals, due to the challenges inherent in expressing and purifying these multi-pass membrane proteins, only a handful of mammalian PATs have been investigated *in vitro*: the palmitoylation of Gα_i_ by DHHC2 and DHHC3, the palmitoylation of PSD-95 by DHHC15, and the autopalmitoylation of DHHC9 and DHHC20^15–19^. These studies often rely on the use of [^3^H]-palmitic acid to quantify the palmitoylated substrate, requiring long exposure times and the hazards of radioactive materials, or else are limited to the kinetics of the autopalmitoylation reaction using enzyme-linked assays or palmitoylation targets of peptide length. We here developed the On-Plate Palmitoylation Assay (OPPA) method as a rapid, click chemistry-based approach to quantifying an intact palmitoylated substrate over time.

The understanding of palmitoylated SNARE proteins has generally been restricted to those lacking transmembrane domains (TMDs), such as SNAP-25 and the vesicular SNARE Ykt6, where the addition of hydrophobic palmitate assists in membrane recruitment^20^. However, the vesicular synaptobrevin-family SNARE protein VAMP7, despite featuring a single TMD at its C-terminus, has been reported to be palmitoylated in a large number of different mammalian global palmitome studies^21–30^. The observation that many of the proteins found to be palmitoylated are in fact integral membrane proteins (up to ∼50-60% of proteins enriched by ABE) suggests the mechanism of regulated palmitoylation is subtler than the simpler model of membrane recruitment^31^. Possible reasons why a transmembrane protein might be palmitoylated include the mediation of protein-protein interactions within the membrane, the induction of conformational changes (e.g. “tilting” within the membrane or the recruitment of soluble tails to the membrane), the driving of localization to liquid-ordered microdomains within the membrane, or as an interplay with other post-translation modifications like ubiquitin ^32^. The palmitoylation of VAMP7 might represent a new mode of transmembrane SNARE protein palmitoylation beyond the relatively straightforward membrane recruitment seen in the soluble SNARE proteins, allowing an additional layer of subtlety to its regulation.

## Results and Discussion

### The composition of the T-cell palmitome changes upon TCR stimulation

Using the acyl-biotin exchange (ABE) method with SILAC-labeled Jurkat T cell lysates allowed us to quantify the T-cell palmitome via quantitative LC-MS/MS (Fig. 1A)^31^. We previously reported the ABE-enriched Jurkat palmitome under resting conditions^31^; here we re-analyze these data with a volcano plot-based approach, allowing us to identify 115 high-confidence palmitoylation candidates, including well-established palmitoylated proteins like Lck and LAT (Fig. 1B). This allows for the direct comparison to the Jurkat palmitome 10 minutes following anti-CD3/anti-CD28 costimulation (Fig. 1C and D, Supp. Fig. 1). After stimulation, 169 proteins are found in the high-confidence palmitoylated pool. Three criteria were used for validation (Fig. 1E): number of previous reports in 13 human palmitome studies, number of predicted palmitoylation sites (CSSPalm 4.0), and number of predicted transmembrane domains (TMDs) (TMHMM 2.0); we see an overrepresentation of all three criteria in our enriched pools versus background unenriched proteins, confirming the enrichment^27,28,39–43,30,31,33–38^.

**Figure 1.**
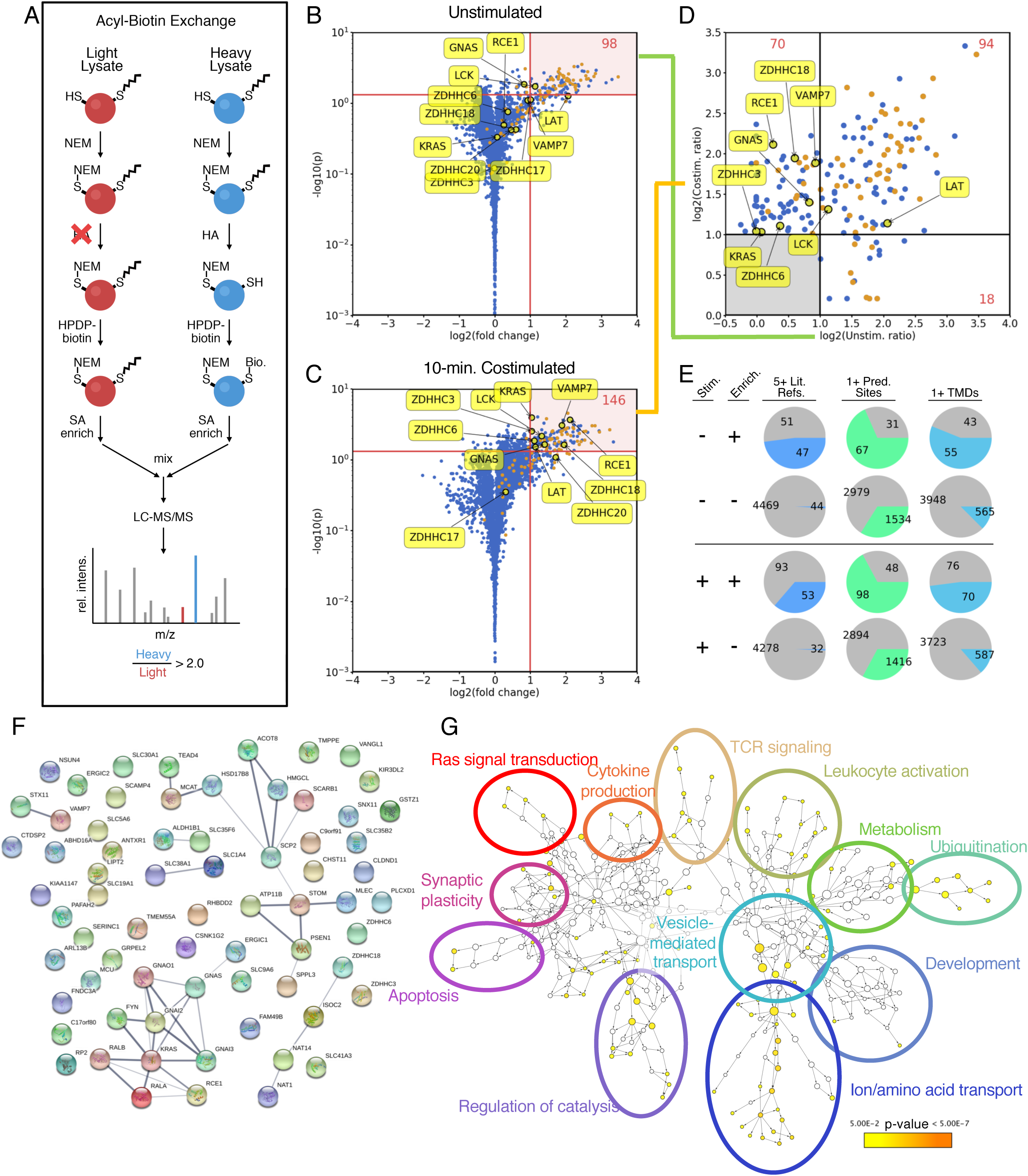
Enrichment of Jurkat T-cell palmitome before and after anti-CD28/anti-CD3-costimulation by acyl-biotin exchange. **A.** Schematic representation of acyl-biotin exchange (ABE) protocol coupled with SILAC for quantitation of palmitoylated proteins via LC-MS/MS. Briefly, Jurkat cells grown in heavy SILAC media (blue) were lysed and free thiols were blocked with NEM. After cleaving palmitoyl groups with HA, previously palmitoylated thiols are biotinylated with HPDP-biotin, allowing for streptavidin enrichment. For control cells grown in light SILAC media (red), HA is omitted, preventing biotinylation. Proteins quantified with heavy/light intensity ratios > 2.0 are considered enriched and palmitoylated. **B.** Volcano plot showing high-confidence palmitoylated proteins prior to stimulation (n=6, p < 0.05). Proteins found in five or more other palmitome studies are highlighted in orange. **C.** Volcano plot showing high-confidence palmitoylated proteins 10 minutes after anti-CD28/anti-CD3 costimulation (n=4, p < 0.05). **D.** Comparison of high-confidence palmitoylated pools before and after stimulation. **E.** Analysis of palmitoylated proteins compared to unenriched (background) proteins, with three metrics: at least five references in previous palmitome studies, at least one predicted palmitoylation site (CSSPalm 4.0, high cutoff), and at least one predicted transmembrane domain (TMHMM 2.0). **F.** STRING functional analysis of proteins enriched for palmitoylation 10 minutes after TCR stimulation. A clear cluster of Gα and Ras-related proteins is observed. Several interesting candidates are highlighted, including VAMP7 and DHHC18, which were investigated in more detail in this study. **G.** Gene Ontology (GO) term enrichment of stimulation-dependent palmitoylation candidates, generated using BiNGO. Cluster size indicates the number of genes in each node, while color indicates statistical significance of enrichment.

Directly comparing these two pools of proteins reveals 94 constitutively palmitoylated proteins, 18 stimulation-dependent depalmitoylated proteins, and 70 stimulation-dependent modified proteins (Supp. Table 1). Lck and LAT are constitutively palmitoylated, while among the stimulation-dependent palmitoylated proteins are the v-SNARE protein VAMP7 and the PAT DHHC18. VAMP7 belongs to the synaptobrevin family of v-SNAREs, and has been shown to be crucial for docking vesicular LAT to the IS and downstream signaling events in CD4+ T cells^11^. DHHC18 is a Golgi-localized PAT, and although it has been described to palmitoylate H-Ras^44,45^, little is known about its enzymatic properties or substrate specificity^19^. As autopalmitoylation is the initial step in the PAT enzymatic mechanism^46^, finding this PAT palmitoylated after TCR stimulation presumably represents an activated state of this enzyme, and may hint at an upregulation of specific PAT activity following TCR stimulation.

Of the stimulation-dependent palmitoylation candidates, approximately half are transmembrane proteins. Grouping the protein interaction networks using STRING analysis reveals a large cluster of Gα, Ras, and Ras-related proteins (Fig. 2A). Intriguingly, within this cluster are GNAI3, GNAO1, and GNAI1/2, all of which feature N-terminal myristoylation motifs, while the Gα proteins GNAQ and GNA13, both of which lack a myristoylation motif, are found to be constitutively palmitoylated. This could suggest an interplay between co-translational myristoylation and post-translational, stimulation-dependent palmitoylation, in which myristoylated Gα proteins are preferentially palmitoylated following TCR stimulation. Indeed, the stimulation-dependent palmitoylation targets feature a slight enrichment of proteins containing myristoylation motifs (8.8%) versus constitutively palmitoylated proteins (6.4%), perhaps hinting at a larger trend. Functionally, the stimulation-dependent palmitoylation candidates are involved in a variety of cellular processes (Fig. 2B), notably Ras signal transduction, TCR signaling, and vesicle-mediated transport.

**Figure 2.**
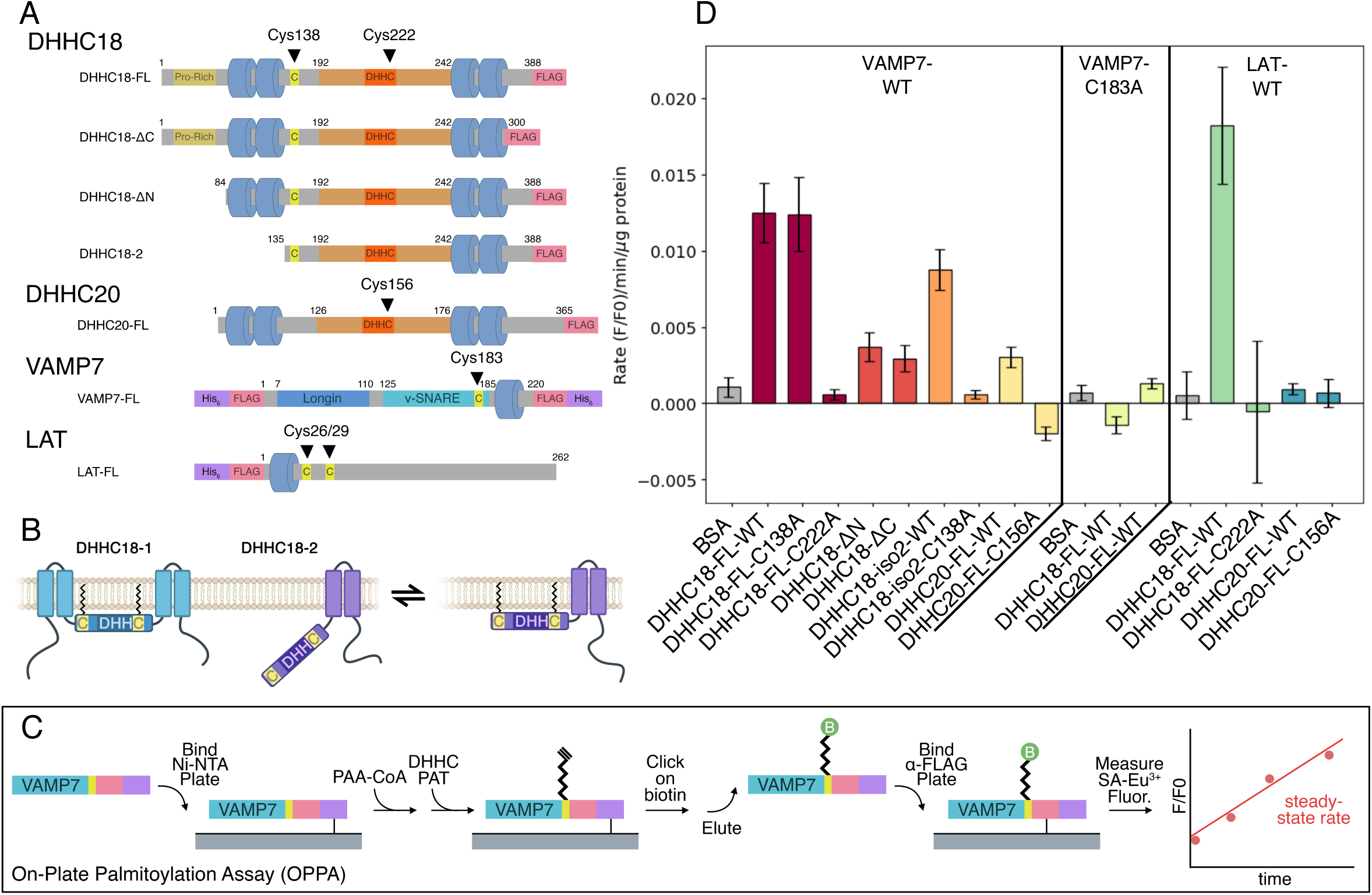
*In vitro* palmitoylation of VAMP7 by DHHC18 and DHHC20. **A.** Schematic cartoon of constructs used, with important cysteines highlighted. Cys222 and Cys156 are the active sites of DHHC18 and DHHC20, respectively. Cys183 in VAMP7 is the site of palmitoylation. **B.** Cartoon model of potential membrane docking of DHHC18 isoform 2 via Cys138. The full-length DHHC18 features the two N-terminal TMDs, securing the DHHC active site to the membrane. The short isoform, lacking these TMDs, may be anchored to the membrane via palmitoylation of Cys138, allowing its active site access to the membrane environment required for its activity. **C.** Schematic representation of the on-plate palmitoylation assay (OPPA). Briefly, the target (VAMP7, featuring both His_6_ and FLAG tags) is bound to a Ni-NTA plate. The DHHC PAT is added with clickable PAA-CoA and the reaction is quenched at specific time points using NEM. Biotin is subsequently clicked on to the PAA-modified target, VAMP7 is eluted and then bound to a anti-FLAG plate. Streptavidin-Eu^3+^ can then be used to quantify palmitoylated VAMP7, and a steady-state slope can be calculated. **D.** Results of OPPA experiments of VAMP7-WT and VAMP7-C183A, palmitoylated by constructs of DHHC18 and DHHC20 (n ≥ 12, geometric mean +/- SEM shown).

### The OPPA method allows for rapid determination of steady-state palmitoylation rates

To investigate these stimulation-dependent palmitoylation candidates in a controlled setting, we designed constructs of DHHC18, DHHC20, VAMP7, and LAT for expression in Sf9 insect cells (Fig 2A, Supp. Figs. 4,5). Although VAMP7 has been reported as palmitoylated in eight other human palmitome studies^23,24,27,28,30,38,39,43,47^, the site of its palmitoylation has not been described. Of the six cysteines in its primary sequence, we found three NEM-modified during our ABE enrichment (data not shown), suggesting they are not palmitoylated, while two others are located in the vesicular lumen, not accessible to a DHHC PAT’s cytosolic active site; the remaining cysteine, Cys183, which is located in an optimal juxtamembrane position, was therefore chosen for mutational studies as the most likely site of palmitoylation^33^.

Likewise, in DHHC18, we sought to investigate the role the N- and C-terminal tails of DHHC18 might play in the enzyme’s stability or activity by designing ΔN and ΔC domain constructs. Moreover, we designed a construct based on the truncated isoform 2 of DHHC18, which we previously hypothesized was preferentially palmitoylated in T cells^31^. Intriguingly, this isoform lacks the two N-terminal TMDs of the full-length protein, yet features a cysteine (Cys138) only three residues from its new N-terminus, which might represent a palmitoylation anchor to the membrane.

In order to improve upon the shortcomings of current *in vitro* palmitoylation assays, we developed the On-Plate Palmitoylation Assay (OPPA) method (Fig 2B). Briefly, purified substrates (e.g. VAMP7) bearing both His_6_- and FLAG-tags are bound to Ni-NTA plates, to which purified DHHC PATs (lacking His_6_-tags) and clickable palmitic acid alkyne (PAA)-CoA are added. The palmitoylation reaction is quenched at specific time points by N-ethylmaleimide (NEM), which both blocks the site of palmitoylation and the DHHC active-site cysteine. After washing, biotin-azide is clicked onto the palmitoylated substrate, which is subsequently eluted from the Ni-NTA plate and bound to anti-FLAG-coated plates. This allows the detection of the biotinylated substrate via fluorescence of Eu^3+^-streptavidin and the determination of the steady-state rate of palmitoylation.

### Examining DHHC18 and DHHC20 activity by OPPA

Using the VAMP7 and LAT constructs as palmitoylation targets and the DHHC18 and DHHC20 constructs as palmitoylating enzymes allowed us to validate the OPPA method and investigate these reactions *in vitro* (Fig 2C). While autopalmitoylation of VAMP7 is minimal at 20 µM PAA-CoA (Supp. Figs 6-8), the rate of palmitoylation is greatly enhanced by full-length DHHC18. The active-site mutant DHHC18-C222A prevents this palmitoylation. DHHC20 was barely capable of palmitoylating VAMP7, while its active-site mutant C156A is also inactive.

Truncating either the N- or C-terminal tails of DHHC18 results in a significant decrease in the palmitoylation rate of VAMP7. Whether this speaks for a role in these cytosolic tails in substrate recognition or engagement, in its enzymatic mechanism, or in simply stabilizing the enzyme will require further investigation. The short isoform DHHC18-2, lacking the N-terminal TMDs, is capable of palmitoylating VAMP7, albeit at a slightly decreased rate. Intriguingly, mutating the proposed palmitoylation anchor cysteine C138 to alanine prevents DHHC18-2’s palmitoylation of VAMP7, while this mutation in full-length DHHC18-1 has no effect; one potential model to explain this is shown in Figure 2B.

Finally, LAT was used as a second palmitoylation target. Full-length DHHC18 was capable of palmitoylating LAT, while DHHC20 failed to do so. This may speak to a specificity for LAT palmitoylation by DHHC18 and not DHHC20. As DHHC20 is located at the PM and turnover of LAT palmitoylation is not expected to be rapid, the specificity of Golgi-localized DHHC18 could perhaps reflect a biological reality; this should be confirmed in a cellular context, perhaps with PAT-specific knockouts and subsequent palmitome enrichment, as was recently performed for DHHC13^45,48^.

### VAMP7 is palmitoylated at Cys183 *in vitro* and *in vivo and is required for Golgi localization*

The proposed palmitoylation site in VAMP7, Cys183, was also mutated to alanine. This prevented its palmitoylation by both DHHC18 and DHHC20, suggesting that this is indeed the primary site of *in vitro* palmitoylation. To confirm this *in vivo*, CRISPR/Cas9 knockout Jurkat cell lines with lentivirally transduced stable re-expression of VAMP7-WT and -C183A constructs were engineered (Supp. Fig. 10). Once generated, ABE was used to enrich all palmitoylated proteins in these cell lines. VAMP7 was found to be palmitoylated in the Jurkat E6.1 and VAMP7-WT re-expression cell lines, but not in the VAMP7 knockout or VAMP7-C183A re-expression cell lines, confirming that Cys183 is the site of VAMP7 palmitoylation in Jurkat T cells (Supp. Fig. 11).

The VAMP7-KO Jurkat cell lines stably expressing VAMP7-WT and VAMP7-C183A were analyzed by TIRFM to observe localization of these two constructs. The re-expressed VAMP7-WT, which is capable of being palmitoylated, is largely found in the Golgi of the VAMP7 knockout/re-expression Jurkat cells, in agreement with earlier observations (Fig 4A)^49^. VAMP7-C183A, however, does not co-localize with the Golgi marker giantin, instead localizing to the cytosol, presumably within vesicles (Fig 3). This supports the conclusion that palmitoylation of VAMP7 at Cys183 acts as a sorting signal for the Golgi^50,51^. Mutating Cys183 prevents this localization, allowing a larger population of VAMP7 to localize to vesicles ^11,52^.

**Figure 3.**
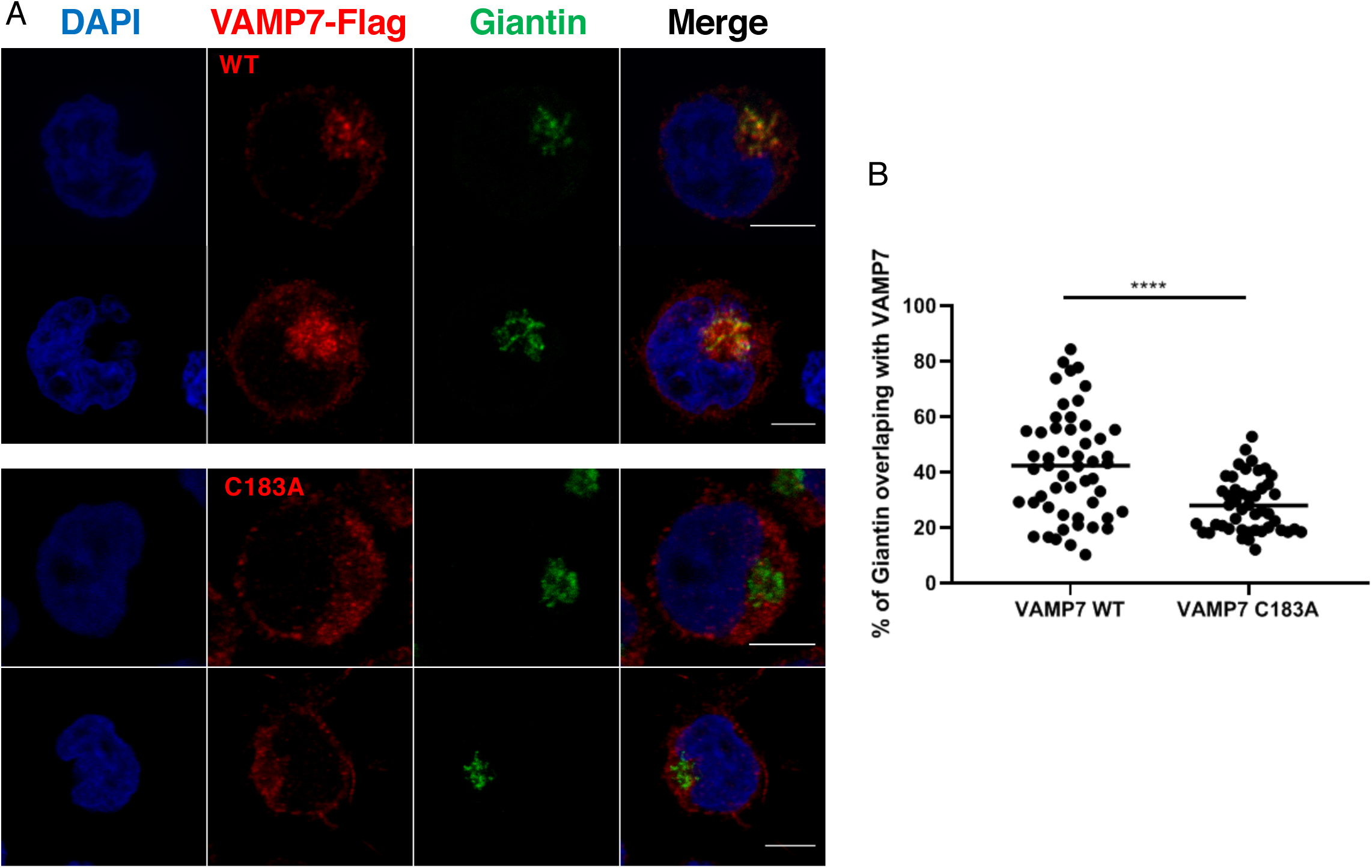
The VAMP7-C183A mutant fails to properly localize to the Golgi. **A** Confocal images showing the relative localization of VAMP7 (red) labelled with anti-Flag Abs and Giantin (green) in VAMP7-KO Jurkat cells expressing the WT VAMP7-Flagged or the mutant VAMP7C183A-Flagged proteins. Scale bar 5 μm. **B** Dot plots showing the quantification of the percentage of Giantin (Golgi marker) overlapping with the VAMP7-flagged proteins (Manderson coefficient). Each dot = one cell; horizontal lines = median. ****P < 0.0001 (one-way ANOVA).

Since VAMP7 has been shown to control membrane trafficking from and to the Golgi^53^, mislocalization of the mutant form of VAMP7 may have some consequences on the trafficking of cargoes in T lymphocytes. It might also alter some cell functions such as polarized transport of proteins or cell migration. The fact that, as shown here, localization of this key v-SNARE protein is regulated upon TCR triggering reinforces the link that exists between TCR-induced signaling and intracellular trafficking^54^.

## Materials and Methods

### Jurkat Cell Culture and Stimulation

Jurkat T cells (clone E6-1) were grown in RPMI 1640 SILAC medium with 10% FCS at 37 °C and 5% CO_2_ (SILAC Quantification Kit, Pierce). Cells were stimulated using anti-CD3 IgM (C305) and anti-CD28 IgM hybridoma supernatants in RPMI without FCS at 37 °C; stimulation was quenched after 10 minutes by the addition of ice-cold PBS, centrifugation (300xg, 5 min, 4 °C), and subsequent shock-freezing of the cell pellets.

### Generation of VAMP7 Knockout in Jurkat Cells

For the generation of sgRNAs the Atum online tool was used (https://www.dna20.com/eCommerce/cas9/input). The sgRNA oligos were then cloned into the pSpCas9(BB)-2A-GFP (PX458) vector (a kind gift from Feng Zhang [Addgene plasmid # 48138; http://n2t.net/addgene:48138; RRID:Addgene_48138]). Jurkat T cells (clone E6-1) were passaged to 0.125 Mio cells/ml the day before transfection. 5 Mio cells were resuspended in 100µl OptiMem with 10 µg pSpCas9(BB)-2A-GFP with the sgRNA CACCGCACACCA-AGCATGTTTGGCA and electroplated with a NEPA21 electroporator (Nepagene) with the following settings: 0.2 mm gap electroporation cuvettes, poring pulse: 175 V, length: 5 ms, interval: 50 ms, number: 2, D. rate: 10%, polarity: +. After 48 hours single clones were sorted by FACS for EGFP into 96-well plates. After the cells had recovered and grown the knockout was validated by Western blotting and sequencing.

### Lentiviral Transduction of Jurkat Cells

Once the VAMP7 knockout cell line was validated, the re-expression of VAMP7 constructs was performed. Wildtype VAMP7 and mutant VAMP7-C183A, both featuring FLAG tags, were cloned into the lentiviral vector LegoiG2 (a kind gift from Boris Fehse [Addgene plasmid # 27341; http://n2t.net/addgene:27341; RRID:Addgene_27341])^55^. HEK293 cells were transfected with the constructs FLAG-VAMP7-WT-LegoiG2 or FLAG-VAMP7-C183A-LegoiG2 plus the third-generation packaging plasmids pMDLg/pRRE with pRSV-Rev and phCMV-VSV-G using lipofectamine. Supernatants containing viruses were harvested after 48 hours and concentrated by centrifugation at 10,000xg and 4°C for 4 to 6 hours in glass tubes. VAMP7 knockout Jurkat cells were transduced with the viruses by spinoculation, as described previously^56^. Cells were resuspended in lentiviral supernatant supplemented with Polybrene (6µg/ml) and spun for 90 min at 37°C at a speed of 800xg. After 72 hours the cells were sorted by EGFP expression via FACS. One week later the cells were sorted again for EGFP expression and also for the expression of CD3 and CD28 surface markers.

### Construct Design and Cloning

Primary sequences of VAMP7, LAT, DHHC18 and DHHC20 were obtained from UniProt. Full-length constructs of VAMP7 and LAT were engineered bearing His_6_- and FLAG-tags, while DHHC18 and DHHC20 were designed with FLAG tags alone, and ordered from Geneart. Truncated constructs of DHHC18 were generated by amplifying FLAG-tagged inserts via PCR. Mutagenesis was performed using the QuikChange Site-Directed Mutagenesis system (Agilent). Constructs were then cloned into the pFastBac1 vector of the Bac-to-Bac system (Thermo Fisher).

### Insect Cell Expression and Purification

Sf9 insect cells were grown in Sf-900 II medium (Thermo Fisher) at 27 °C. Baculoviruses were generated by transfecting insect cells with constructs in the pFastBac1 vector according to the Bac-to-Bac protocol. P4 viral stocks were then used to infect larger cultures, and expressed proteins were harvested 3-5 days following infection (∼70-80% cell viability). Cell pellets were shock frozen and lysed in lysis buffer containing 50 mM NaH_2_PO_4_ (pH 7.2) (Roth), 300 mM NaCl (Roth), 0.2% Triton X-100 (Roth), 1 mM phenyl methanesulfonyl fluoride (PMSF) (Sigma), cOmplete protease inhibitor cocktail (Roche), and DNase (New England BioLabs) and RNase (Pancreac AppliChem) on ice for 30 minutes. Lysates were cleared by centrifugation (16,000xg, 4 °C, 10 min.) and supernatants were loaded onto anti-FLAG M2 affinity gel (Sigma) at 4 °C. After washing, proteins were eluted using 0.1 M glycine at pH 3.5 and concentrated using Vivaspin spin filters (VWR) with appropriate MW cut-offs. After concentration, 0.08% n-dodecyl β-D-maltoside (DDM) (Sigma) and cOmplete protease inhibitor cocktail was added and constructs were dialyzed overnight at 4 °C into 50 mM NaH_2_PO_4_ (pH 6.5) and 10% glycerol (Roth). Protein concentration was determined by BCA assay (Pierce) and expression was confirmed by anti-FLAG Western blot. Purified PATs were extremely sensitive to environmental conditions, precipitating at concentrations greater than ∼5-10 µM and losing activity after freezing or 3-5 days storage at 4 °C. For this reason, all purified PATs were used immediately for the OPPA assays following dialysis.

### Western Blotting

To confirm the stimulation via phosphotyrosine and phospho-ERK1/2, and to confirm expression/purification of constructs expressed in insect cells, lysates were run on SDS-PAGE gels and transferred to nitrocellulose membranes using standard methods. Antibodies used for detection: 4G10 Platinum anti-phosphotyrosine (Millipore), anti-pERK (E-4) (Santa Cruz), anti-β-actin (AC-15) (Sigma Aldrich), anti-LAT (Upstate (Millipore)) and anti-FLAG-HRP (Sigma Aldrich). The self-produced VAMP7 antibody was a kind gift of Dr. Andrew Peden (The University of Sheffield, UK).

### Acyl Biotin-Exchange

The ABE protocol was followed as described previously^31,57^. Briefly, stimulated Jurkat SILAC pellets were lysed in lysis buffer containing 50 mM Tris (pH 7.4) (Roth), 150 mM NaCl (Roth), 10 mM MgCl_2_ (AppliChem), 10mM KCl (Sigma-Aldrich), 500 µM EDTA (Roth), 100 µM Na_3_VO_4_ (Sigma-Aldrich), 20 mM NEM (Thermo), 1 mM PMSF, 1.7% Triton X-100, 1 mM tris(2-carboxyethyl)phosphine (TCEP) (Sigma-Aldrich), 100 µM methyl arachidonyl fluorophosphonate (MAFP) (Sigma-Aldrich), and cOmplete protease inhibitor cocktail (PI-cocktail) on ice for 30 minutes. Lysates were cleared by centrifugation (16,100xg, 4 °C, 10 min.). Chloroform-methanol (CM) precipitations were used between each chemical labeling step to ensure complete remove of unreacted reagents. The cleared lysates were subjected to a CM precipitation, then resuspended in buffer containing a 50 mM Tris (pH 7.4), 5 mM EDTA, 1% SDS (Roth), 125 mM NaCl, 1 mM PMSF, 0.2% Triton X-100, 20 mM NEM, and PI-cocktail and incubated overnight at 4 °C. NEM was then removed by three sequential CM precipitation steps, and enriched samples (e.g. heavy lysates) were resuspended in buffer containing 50 mM Tris (pH 7.4), 5 mM EDTA, 125 mM NaCl, 1% SDS, 574 mM hydroxylamine (HA) (Sigma-Aldrich), 820 µM EZ-Link HPDP-biotin (Thermo), 0.2% Triton X-100, 1 mM PMSF and PI-cocktail; control samples (e.g. light lysates) were resuspended in a similar buffer, omitting HA. After incubation for one hour at room temperature, a single CM precipitation removed HA, and samples were biotinylated in a buffer containing 50 mM Tris (pH 7.4), 5 mM EDTA, 125 mM NaCl, 164 µM EZ-Link HPDP-biotin, 0.2% Triton X-100, 1 mM PMSF and PI-cocktail for one hour at room temperature. Samples were then washed with three sequential CM precipitations and resuspended in buffer containing 50 mM Tris (pH 7.4), 5 mM EDTA, 125 mM NaCl, 0.1% SDS, 0.2% Triton X-100, 1 mM PMSF and PI-cocktail. SILAC-labeled heavy and light samples were then mixed and allowed to bind to streptavidin-agarose beads (Novagen) for 90 min at room temperature. After four sequential washings, biotinylated peptides were eluted by cleaving the HPDP-biotin-cysteine disulfide linkage in a buffer containing 1% beta-mercaptoethanol (2-ME) (Roth) at 37 °C for 15 min. Eluates were precipitated via trichloroacetic acid (TCA) precipitation and resuspended in a smaller volume of SDS-PAGE sample buffer containing 0.2% Triton X-100, boiled for 5 min. at 95 °C, then run on a 4-12% Bis-Tris gel (NuPAGE Novex). After Coomassie staining, gels were used for in-gel tryptic digestion and subsequent LC-MS/MS. For the stimulated Jurkat cells, four individual ABE enrichments were performed (n=4).

### Quantitative LC-MS/MS and Proteomic Data Analysis

Coomassie-stained gel lanes from the ABE enrichment were cut into 30 equal-sized bands and in-gel tryptic digest was performed as described previously^31^. Digested peptides were resuspended in 6 µl 0.1% (v/v) TFA and 5% (v/v) acetonitrile per band. Analysis of the peptides was performed via reversed-phase capillary liquid chromatography (Ultimate 3000 nanoLC system (Thermo Scientific)) coupled to an Orbitrap Elite mass spectrometer (Thermo Scientific). LC separations were performed on a capillary column (Acclaim PepMap100, C18, 3 µm, 100 Å, 75 µm i.d.×25cm, Thermo Scientific) at an eluent flow rate of 300 nL/min using a linear gradient of 3–25% B after 53 min with a further increase to 80% B after 80 min. Mobile phase A contained 0.1% formic acid in water, and mobile phase B contained 0.1% formic acid in acetonitrile. Mass spectra were acquired in a data-dependent mode with one MS survey scan with a resolution of 60,000 (Orbitrap Elite) and MS/MS scans of the 15 most intense precursor ions in the linear trap quadrupole were used.

Identification and SILAC quantification were performed using MaxQuant software (version 1.4.1.1) with the Uniprot human protein database (July 2013). Proteins were filtered with the following criteria: i) not identified only by site, reverse, or known contaminant (MaxQuant), ii) at least one unique peptide, and iii) at least two razor + unique peptides. The unstimulated Jurkat ABE enrichment we published earlier^31^ (n=6) was re-analyzed using the following new procedure, allowing direct comparison to the stimulated Jurkat ABE dataset described here (n=4). First, missing values were imputed with the median normalized heavy/light ratio of the specific LC-MS/MS run – in many cases this was very close to 1.0 (representing the background). H/L ratios were then transformed to log2 values and the mean H/L ratio and one-sample t-test p values were calculated. Volcano plots were generated plotting the log2 fold-change to the –log10(p) value, giving rise to a pool of high-confidence palmitoylated proteins. Enriched pools were analyzed based on three criteria, described previously^31^: i) previous reports in other palmitome studies, ii) predicted palmitoylation sites (CSS-Palm 4.0), and iii) predicted TMDs (TMHMM 2.0)^33,34^.

### On-Plate Palmitoylation Assay (OPPA)

Purified targets (VAMP7, LAT) were diluted to 1 µM in buffer containing 50 mM NaH_2_PO_4_ (pH 6.5), 500 mM NaCl, 1 mM TCEP and 0.08% DDM. For each time point (0, 5, 10 and 20 min), the diluted target was divided into 12 wells (50 µl each) of a 96-well Ni-NTA-coated plate (Pierce) and allowed to bind for 1 hour. Palmitic acid alkyne (PAA)-CoA was generated by preparing 50 mM Tris (pH 8.1), 14 mM MgCl_2_, 0.08% DDM, 0.1 mM EDTA, 1.6 mM ATP (Roth), 400 µM coenzyme A (Sigma), 40 µM PAA (Avanti), 0.05 UN/ml acyl-coenzyme A synthetase (Sigma) and 1% BSA (Sigma) and incubating at 37 °C for 1 hour. This was diluted two-fold (yielding [PAA-CoA] ∼ 20 µM) with 0.3-2 µM purified DHHC PAT (or 1 µM BSA for negative control) with 150 mM NaH_2_PO_4_ (pH 6.5), 300 mM NaCl and 0.08% DDM. Ni-NTA plates were washed 4 times with PBS + 0.05% Tween-20 (Sigma-Aldrich) (PBST) using a plate washer (Tecan) and 100 µl of the enzyme mixture was added. Reactions were carried out at 37 °C with shaking for designated time points, and the reaction was quenched by washing with buffer containing 50 mM NaH_2_PO_4_ (pH 6.5), 300 mM NaCl, 40 mM NEM and 0.1% Triton X-100. After the complete time course, plates were washed 4 times with PBST and click chemistry was performed by the addition of 50 µl buffer containing 50 mM NaH_2_PO_4_ (pH 6.5), 300 mM NaCl, and 1% Triton X-100. Click chemistry reagents were added as described previously^58^; briefly, to each 50 µl, the following reagents were added in this order, with vortexing between each step: 1 µl 1 mM biotin-azide (Thermo Fisher) in DMSO (Sigma), 1 µl 50 mM TCEP, 3 µl 16 mM Tris[(1-benzyl-1*H*-1,2,3-triazol-4-yl)methyl]amine (TBTA) (Sigma-Aldrich) prepared in 20% DMSO and 80% t-butanol (Sigma), and 1 µl 50 mM CuSO_4_ (Sigma). Click chemistry was performed at 37 °C for 3 hours with shaking. After washing the plate 6 times with PBST, palmitoylated targets were eluted with 100 µl buffer containing 50 mM NaH_2_PO_4_ (pH 6.5), 300 mM NaCl, 1% Triton X-100, 1 mM EDTA and 200 mM imidazole (Roth) for 30 min at 37 °C. Eluates were transferred to high-binding plates (Greiner) coated for 3 hr at room temperature with monoclonal anti-FLAG (Sigma-Aldrich) and blocked for 2 hr at room temperature with 3% BSA and allowed to bind overnight at 4 °C with shaking. Plates were then washed 6 times with PBST and buffer containing PBS and 0.2 µg/ml Eu^3+^-streptavidin (PerkinElmer) and incubated at room temperature for 1 hour. After washing 8 times with PBST, fluorescence was measured in 120 µl enhancer solution (15 µM beta-naphthoyltrifluoroacetone (Sigma), 50 µM tri-n-cotylphosphine oxide (Sigma), 6.8 mM potassium hydrogen phthalate (Roth), 100 mM acetic acid (Roth) and 0.1% Triton X-100) using a plate reader (Victor^3^V, PerkinElmer) with excitation and emission wavelengths of 340 nm and 614 nm, respectively. All measured fluorescence values were normalized to both the t=0 fluorescence value and the µg of protein (DHHC PAT or BSA) used in the reaction, and steady-state rates were calculated over the 20-minute reaction time.

### Immunofluorescence

#### Coverslips & dishes preparation

12mmø coverslips (VWR, 631-0666) were pre-coated with 0.02% of poly-L-Lysine for 20 min at room temperature and were washed 3 times in water before being dried and kept for maximum 2 days. 100,000 cells were incubated on coverslips for 30 min in PBS.

#### Fixation

Cells were fixed with 4% paraformaldehyde (Life Technologies, FB002) for 15 min at room temperature, washed once in PBS and excess of paraformaldehyde was quenched for 10 min with PBS 10 mM glycine (Thermo Fisher Scientific, G8898). Coverslips were kept at 4°C in PBS until permeabilization and staining.

#### Staining

Cells were permeabilized for 30 min at room temperature with PBS + 0.2% Bovine Serum Albumin (BSA, Euromedex, 04-100-812) and 0.05% Saponin (Sigma-Aldrich, S4521). Cells were then incubated for 1 hour at room temperature with primary antibody, then washed 3 times with PBS 0.2% BSA 0.05% Saponin and incubated protected from light for 20 min in the same buffer with spun secondary antibodies. After washing once with PBS BSA Saponin, and once with PBS, coverslips were soaked three times in PBS, three times in water, and mounted on slides.

#### Mounting

For regular confocal microscopy, coverslips were mounted with 4-6 µL Fluoromount G (SouthernBiotech, 0100-01) on slides (KNITTEL Starfrost) and dried overnight protected from light before microscope acquisition.

#### Microscope

Images were acquired with a Leica DmI8 inverted microscope equipped with an SP8 confocal unit using either a 40X (1.35NA) or 63X (1.4NA) objective. Single plane images or Z-stack of images were acquired (pixel size around 60 nm).

#### Analysis of VAMP7 colocalization with Giantin

Z-stack (0.5 µm) images of similarly dimensioned Jurkat cells were chosen. In this z-stack, an ROI surrounding the Golgi was defined based on Giantin staining. Within each ROI, masks based on both Giantin and VAMP7 stainings were created by thresholding. Automatic colocalization assays were performed with Mander’s overlap coefficient, using the JACoP plugin for ImageJ64.

#### Antibodies

Anti-Flag (1/100) was from Sigma Aldrich (F3165). Anti-Giantin (1/100) was produced by the recombinant antibody platform of the Institut Curie, Paris, France. Anti–rabbit Ig Alexa Fluor 488 (1/200) and anti–mouse Ig Alexa Fluor 568 (1/200) antibodies were from Thermo Fisher Scientific (A11034 and A11004 respectively).

## Acknowledgements

Special thanks to Frank Kuppler, Ellie Fox, Michael Schümann, and Heike Stephanowitz for technical support. Special thanks to Andrew Peden (The University of Sheffield, UK) for gifting us the anti-VAMP7 antibodies. CF and BB are supported by the DFG (TRR186, Project A05). CH is supported by funds from Institut Curie, Institut National de la Santé et de la Recherche Médicale (INSERM) and ANR (ANR-10-IDEX-0001-02 PSL*, and ANR-11-LABX-0043).

---

**Suppl. Table 1.**
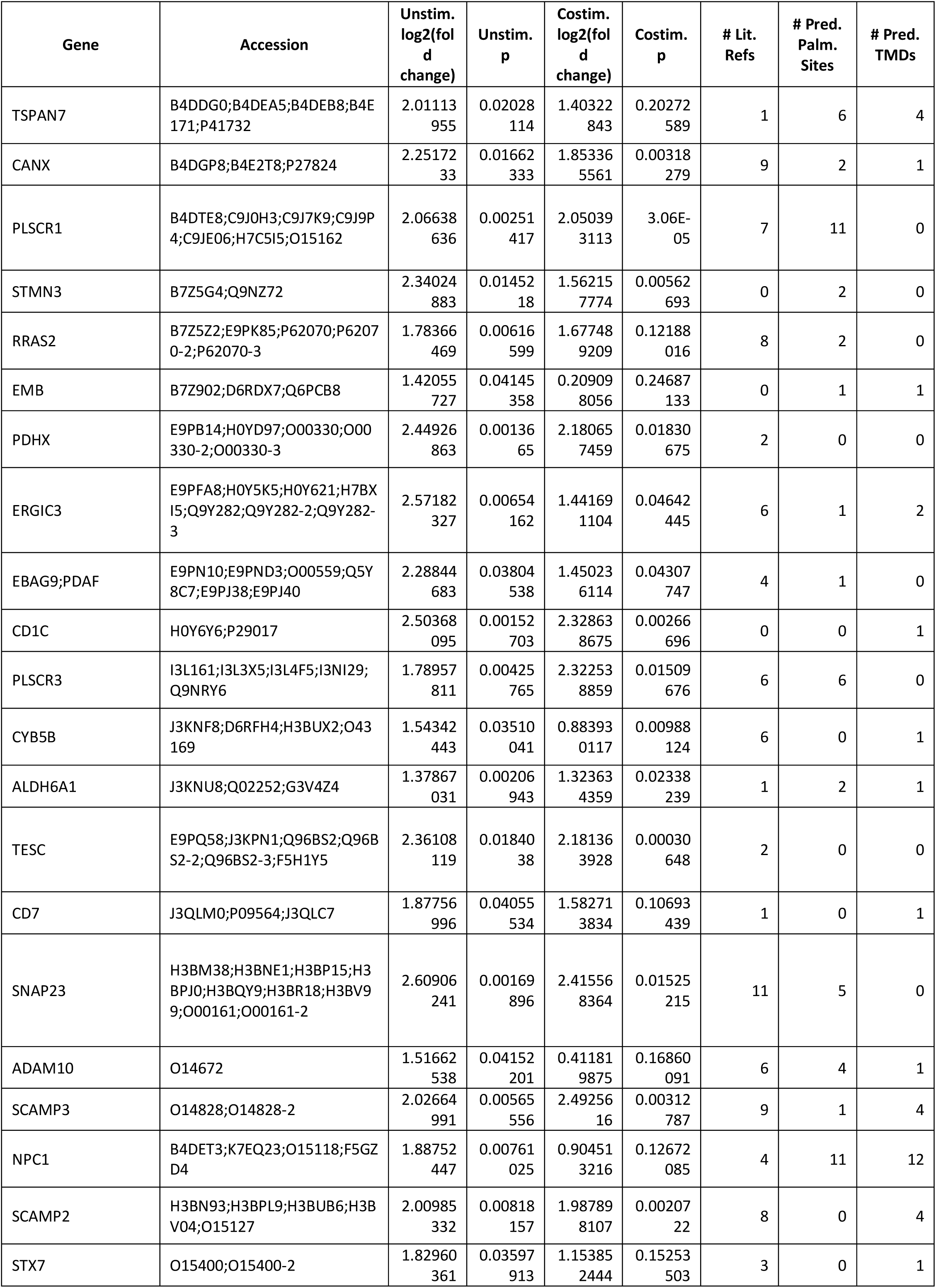
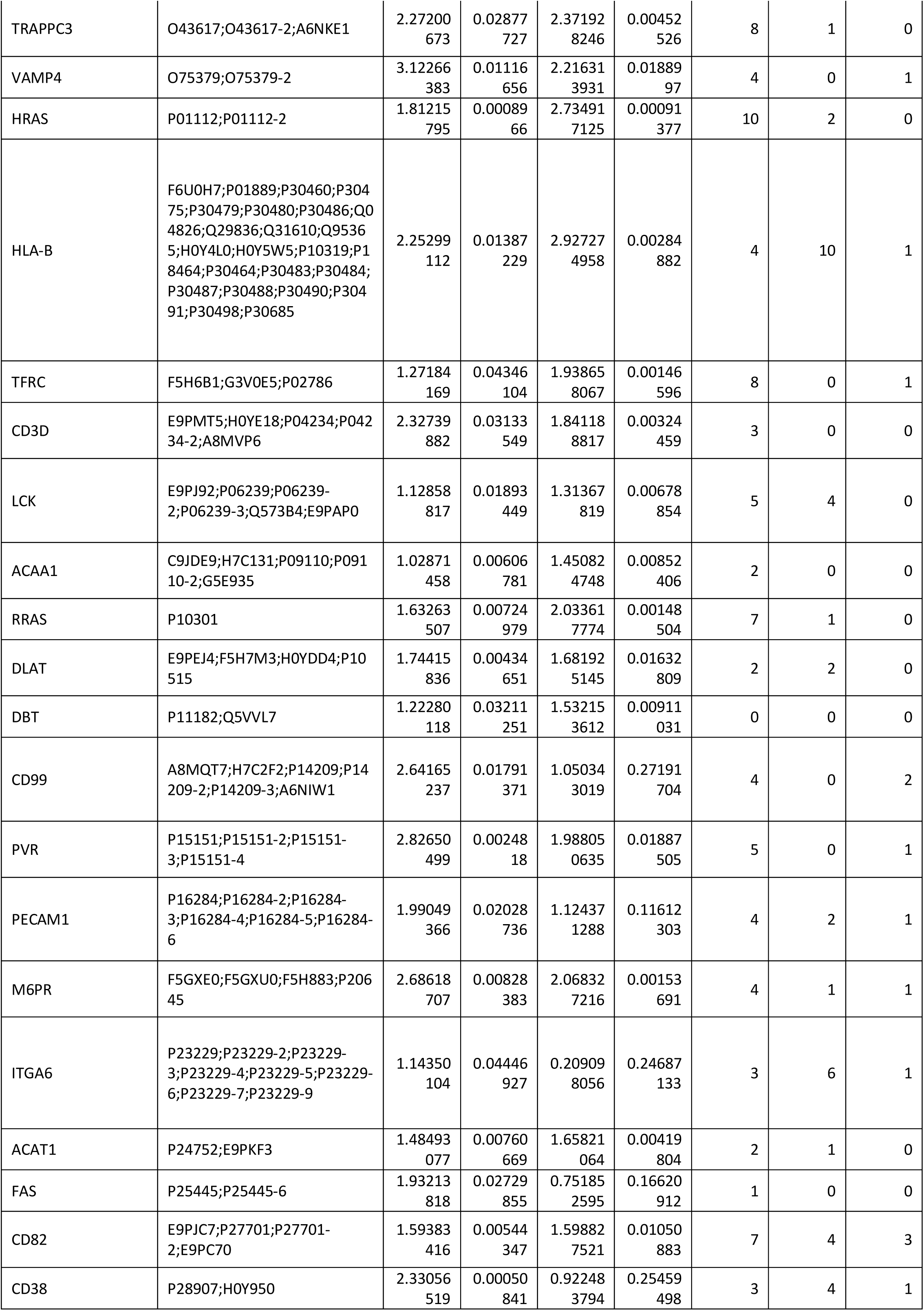
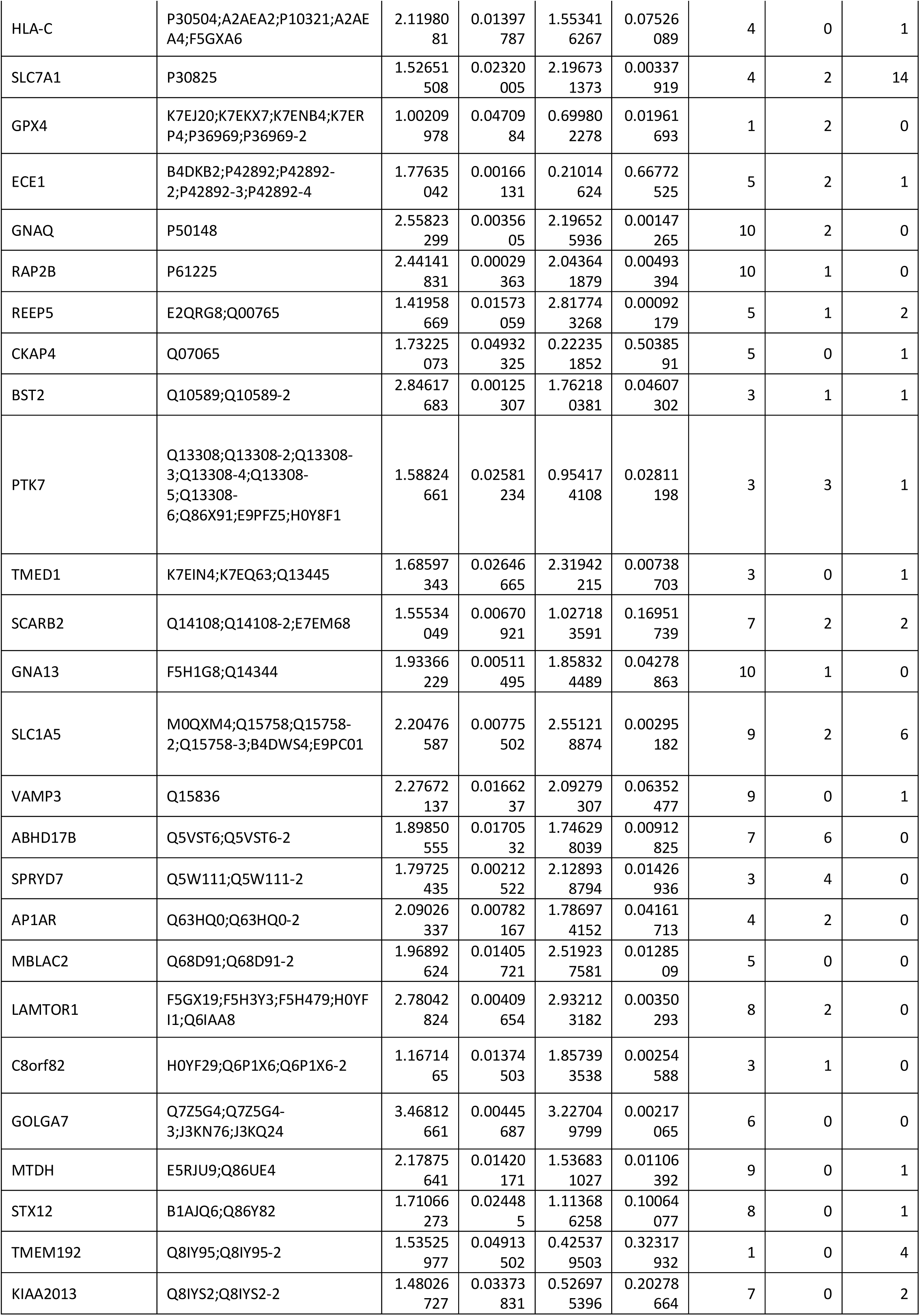
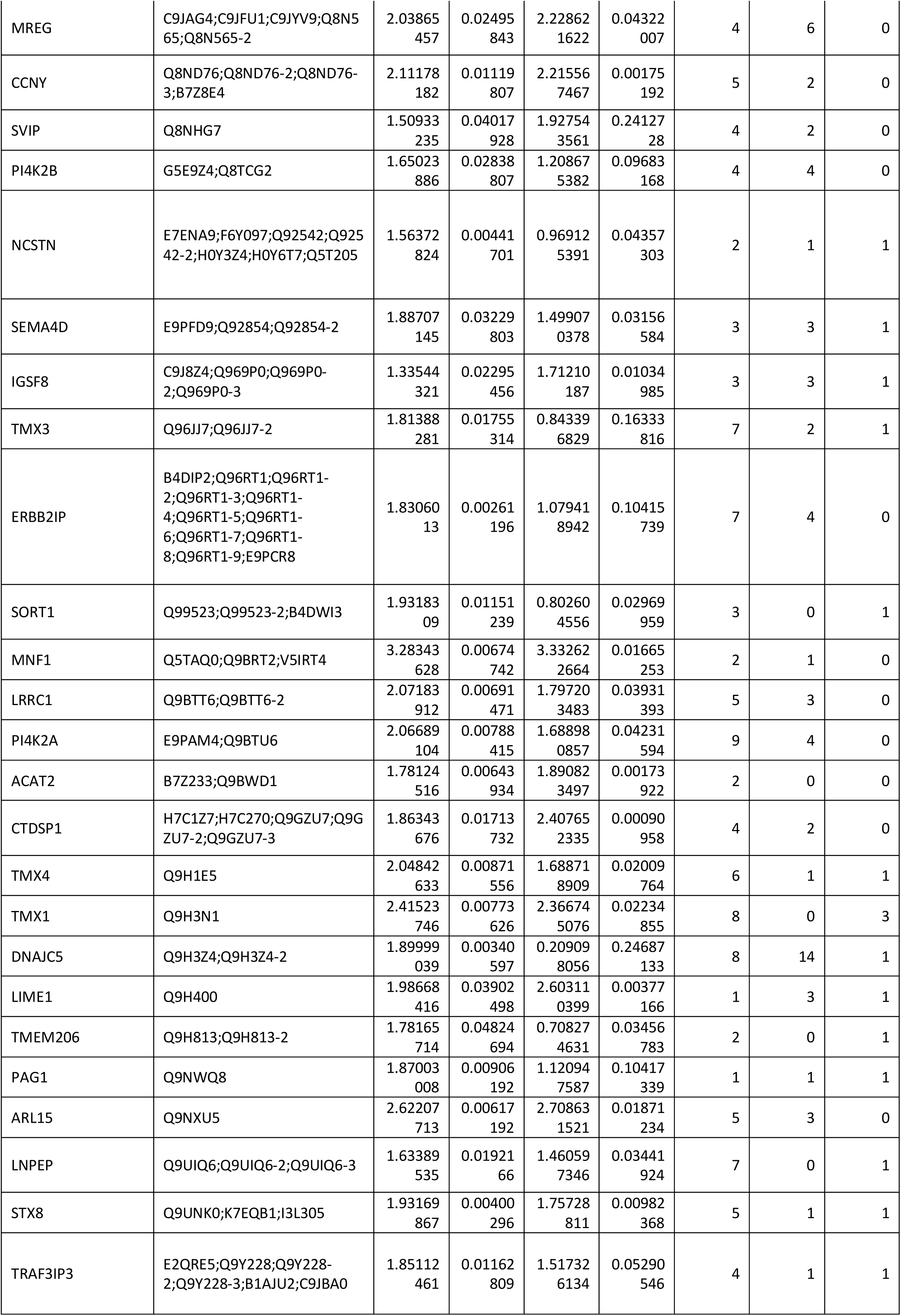
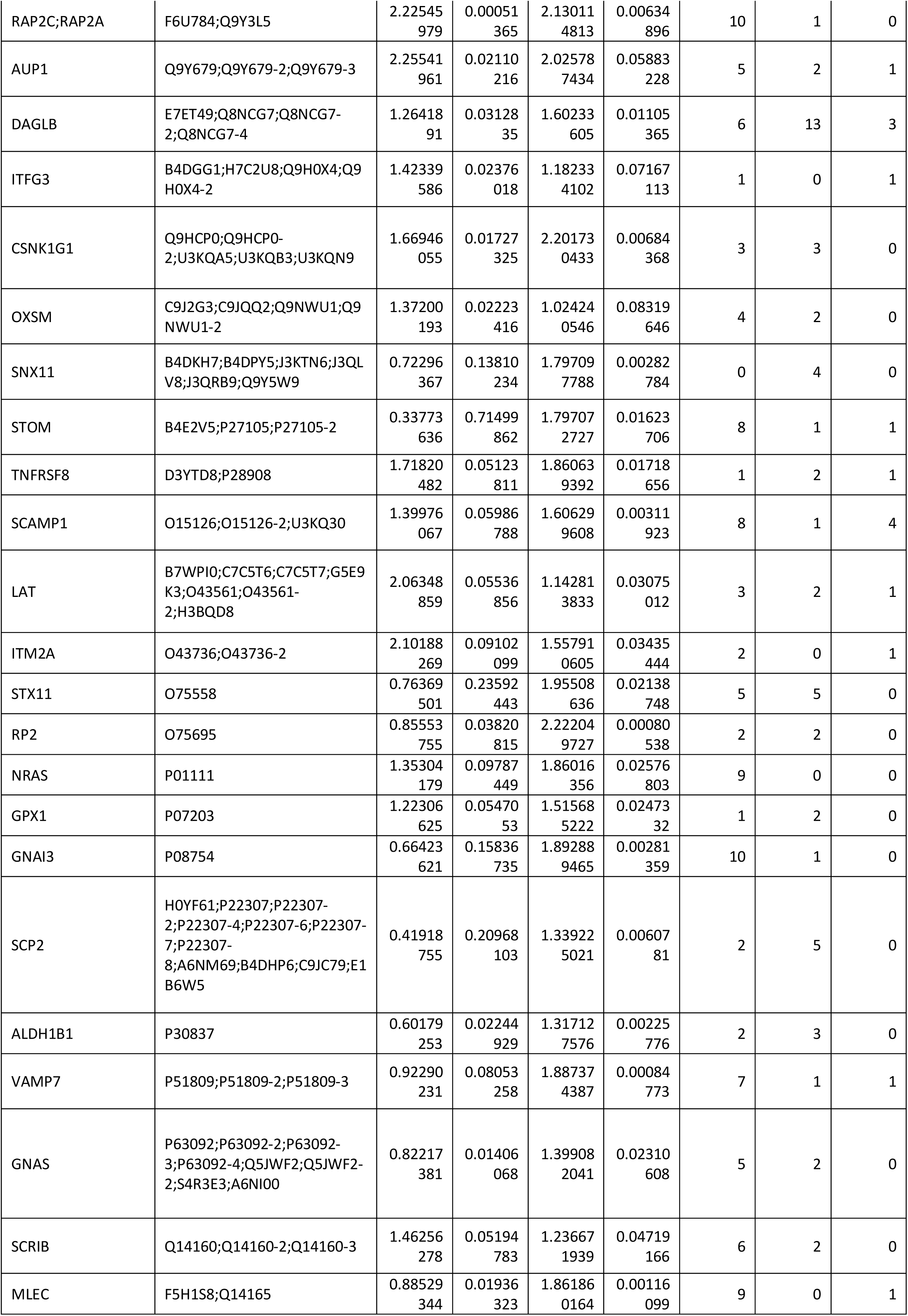
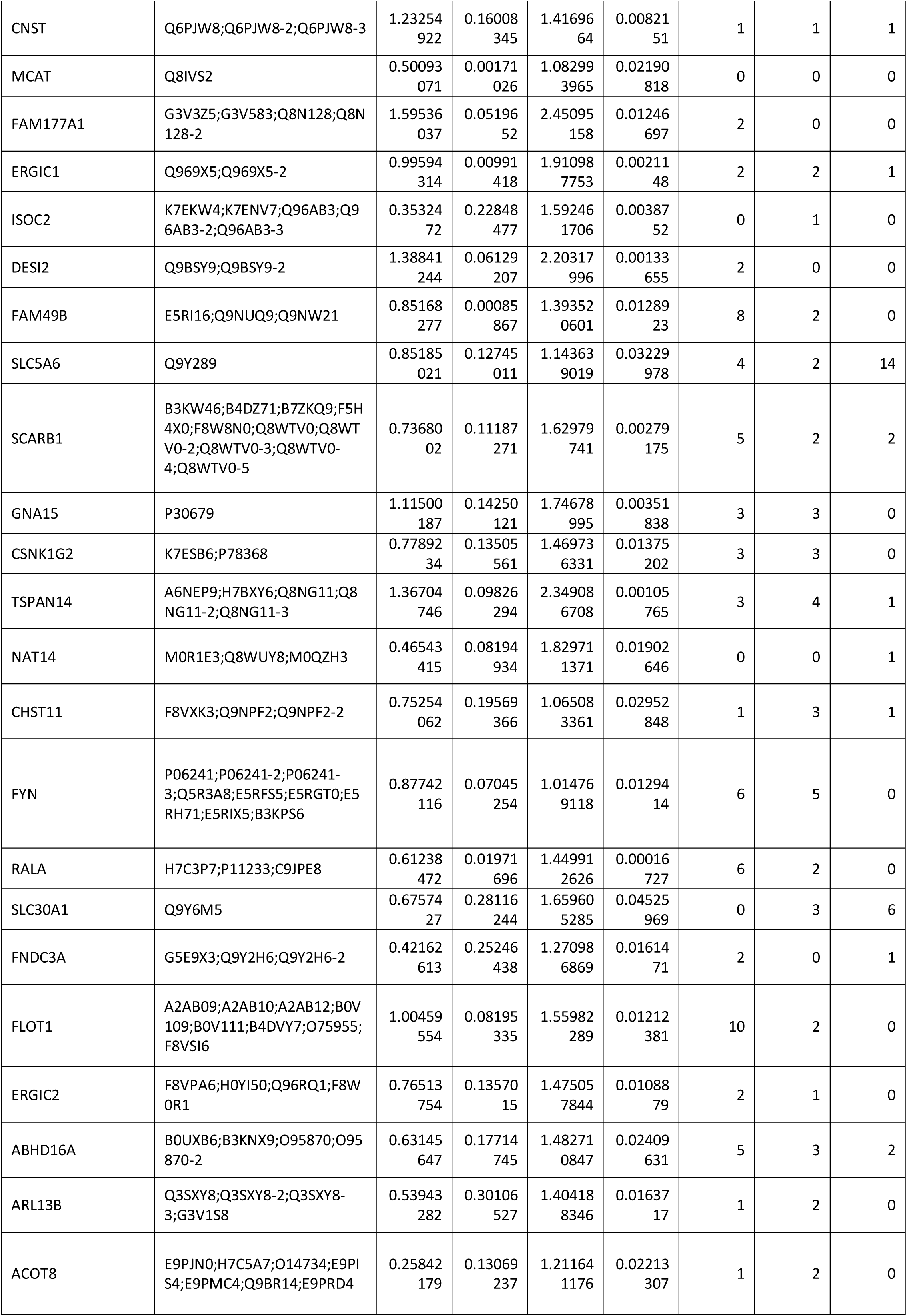
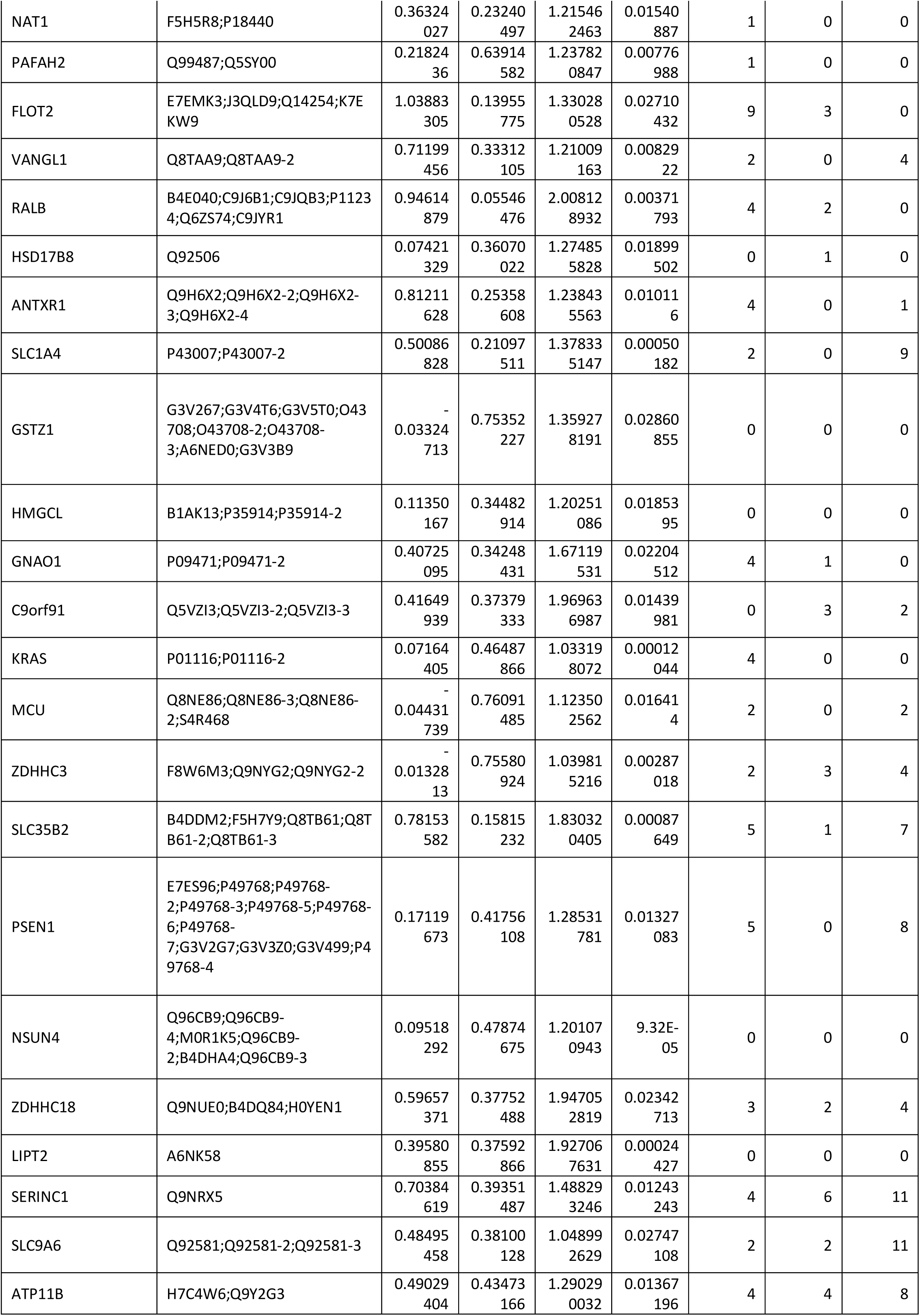
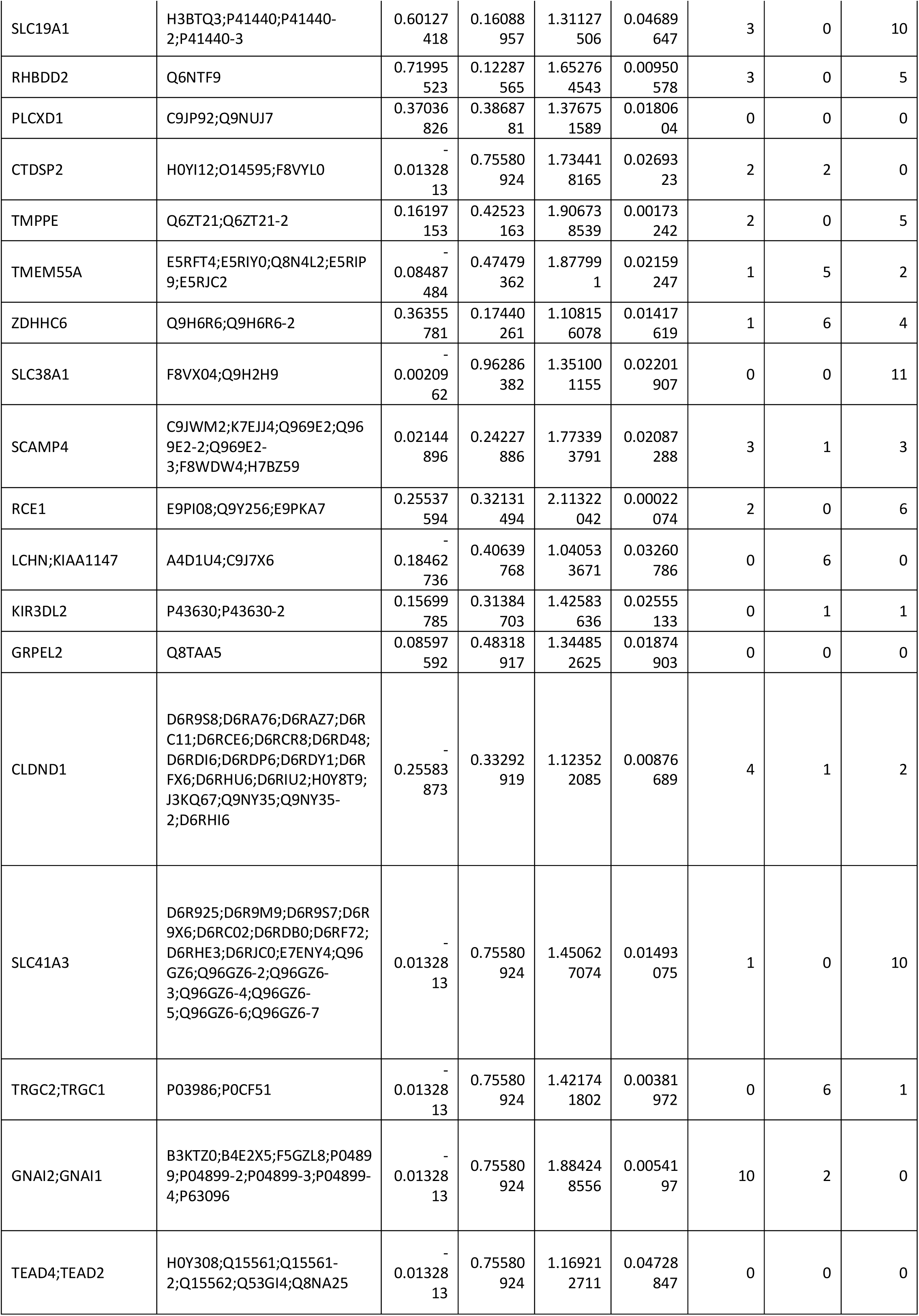
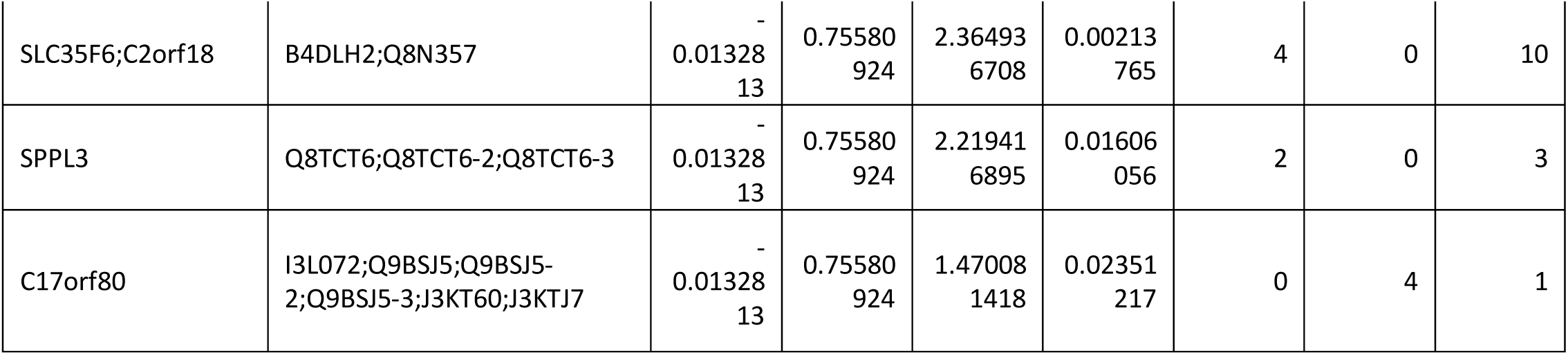
Proteins enriched before and 10 min. after T-cell stimulation

