## Supplementary material for "Dynamic Palmitoylation Events Following T-Cell Receptor Signaling": Supplmental Figures

### Supplemental Figures

Supp. 1

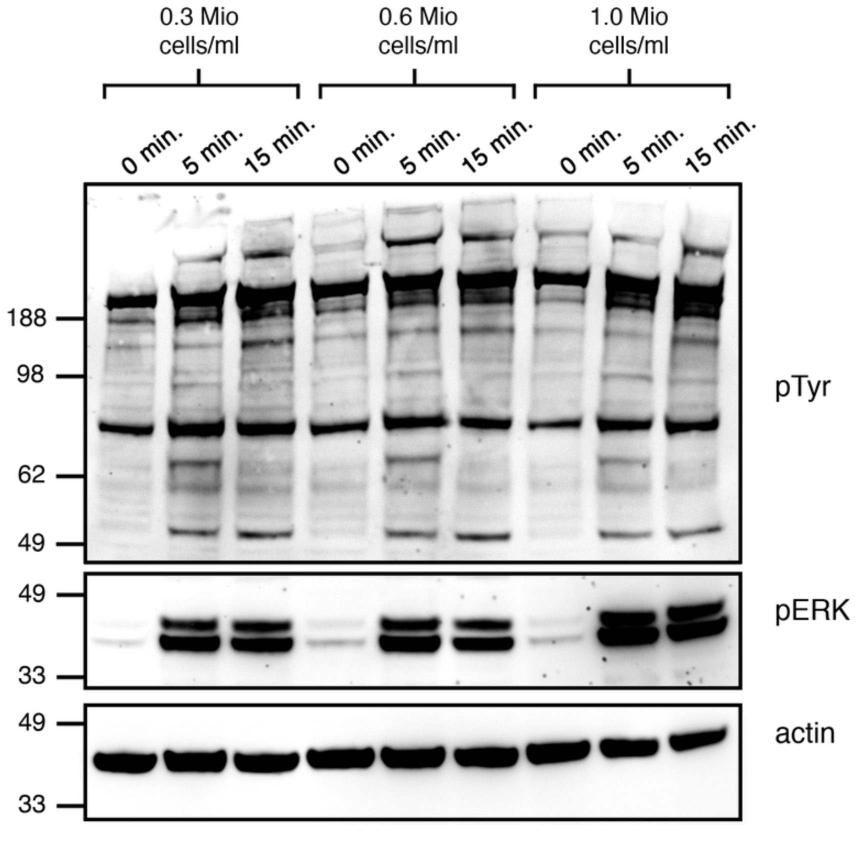


**Supplemental Figure 1.** Western blot showing increased phosphorylation events following anti-CD28/anti-CD3 costimulation of Jurkat T cells at three different cell densities.

Supp. 2

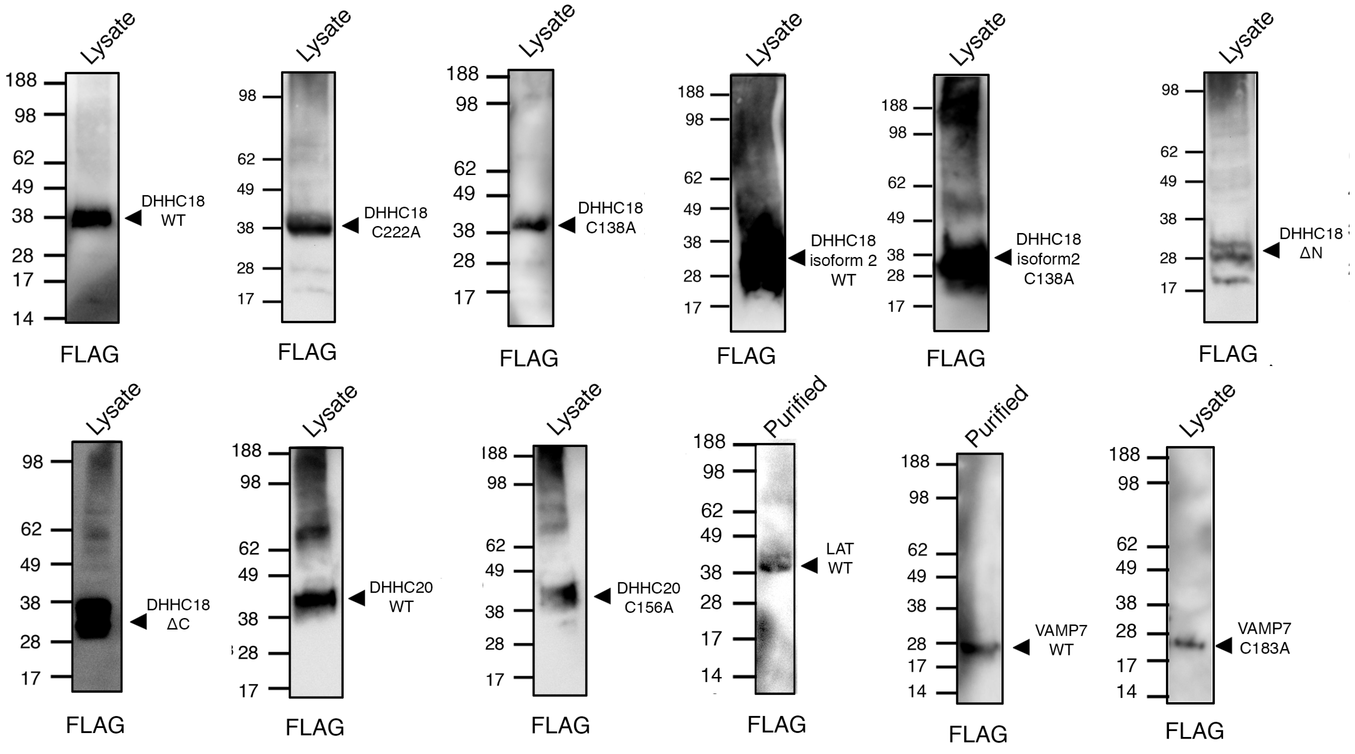


**Supplemental Figure 2.** Western blots showing insect cell-expressed constructs used. Anti-FLAG antibodies were used for all Western blots.

Supp. 3

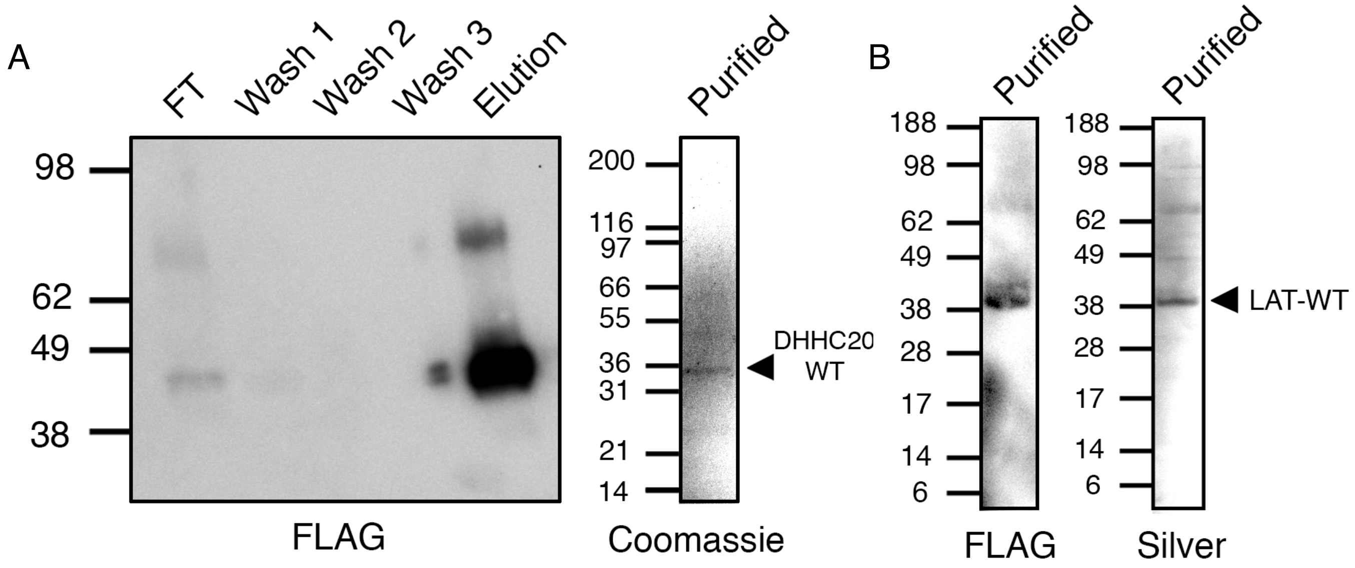


**Supplemental Figure 3. A.** Western blot of FLAG purification of DHHC20-WT. Also shown is a Coomassie-stained SDS-PAGE gel of the purified protein. **B.** Anti-FLAG Western blot and silver stained SDS-PAGE gel of purified target LAT-WT.

Supp. 4

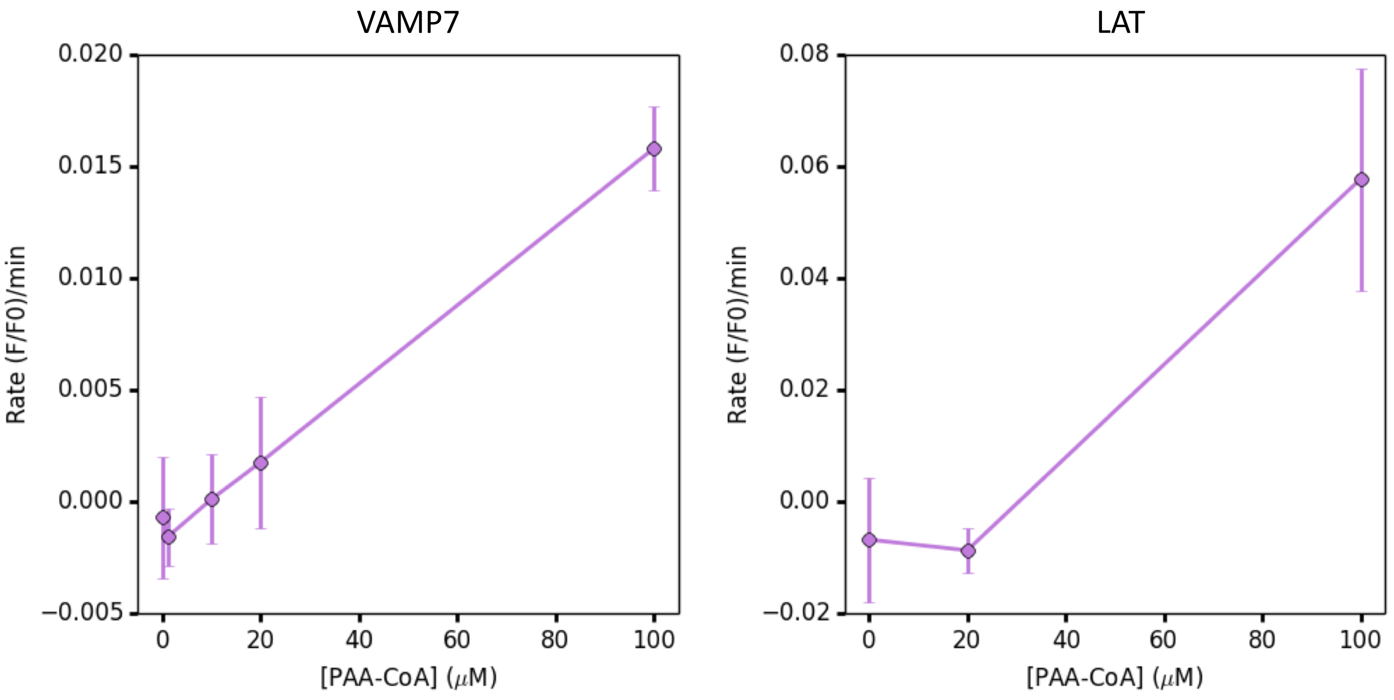


**Supplemental Figure 3.** Titration of PAA-CoA showing autopalmitoylation rates of VAMP7-WT and LAT-WT using the OPPA method.

Supp. 5

**Supplemental Figure 5. A.** Enzyme-linked assay confirming the generation of clickable PAA-CoA. The formation of palmitoyl-CoA from palmitic acid is used as a positive control. **B.** The enzymatic reactions used in (A), allowing the observation of NADH oxidation at A340 as a proxy for PAA-CoA production

A

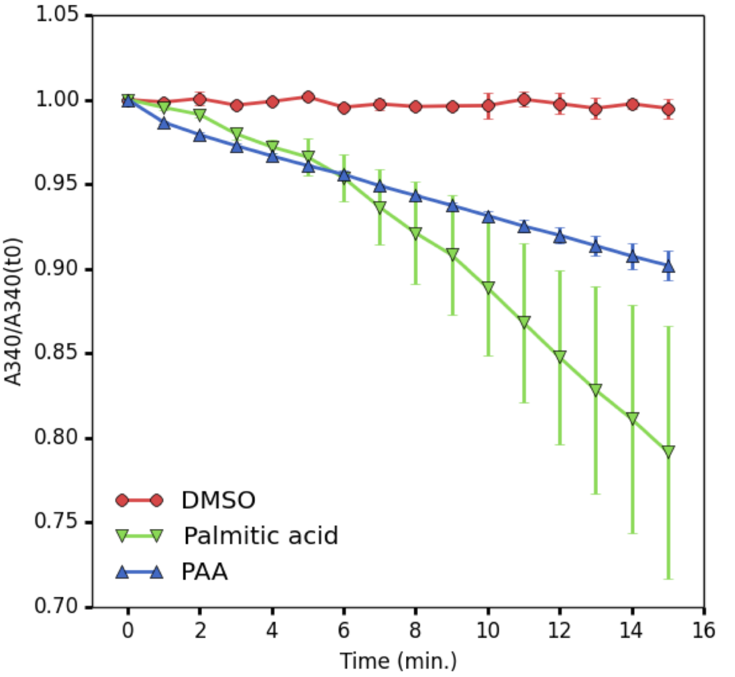


B

$$\text{CoA + PAA + ATP} \xrightarrow{\text{Acyl CoA Synthetase}} \text{PAA-CoA + AMP + PP}_i$$

$$\text{ATP + AMP} \xrightarrow{\text{Myosin Kinase}} \text{2 ADP}$$

$$\text{2 ADP + 2 Phospho(enol)pyruvate} \xrightarrow{\text{Pyruvate Kinase}} \text{2 ATP + 2 Pyruvate}$$

$$\text{2 Pyruvate + 2 NADH} \xrightarrow{\text{L-Lactate Dehydrogenase}} \text{2 L-Lactate + 2 NAD}$$

Supp. 6

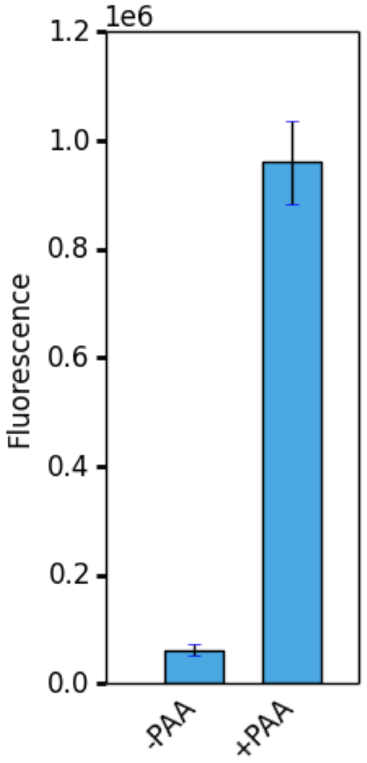


**Supplemental Figure 6.** Confirmation of click chemistry conditions. Jurkat T cells were fed with PAA and incubated for 4 hours at 37 ˚C with 5% CO2. After lysis, biotin-azide was clicked on to all PAA-modified proteins. The lysates were bound to a high-binding plate and $Eu^{3+}$-streptavidin fluorescence was measured.

Supp. 7

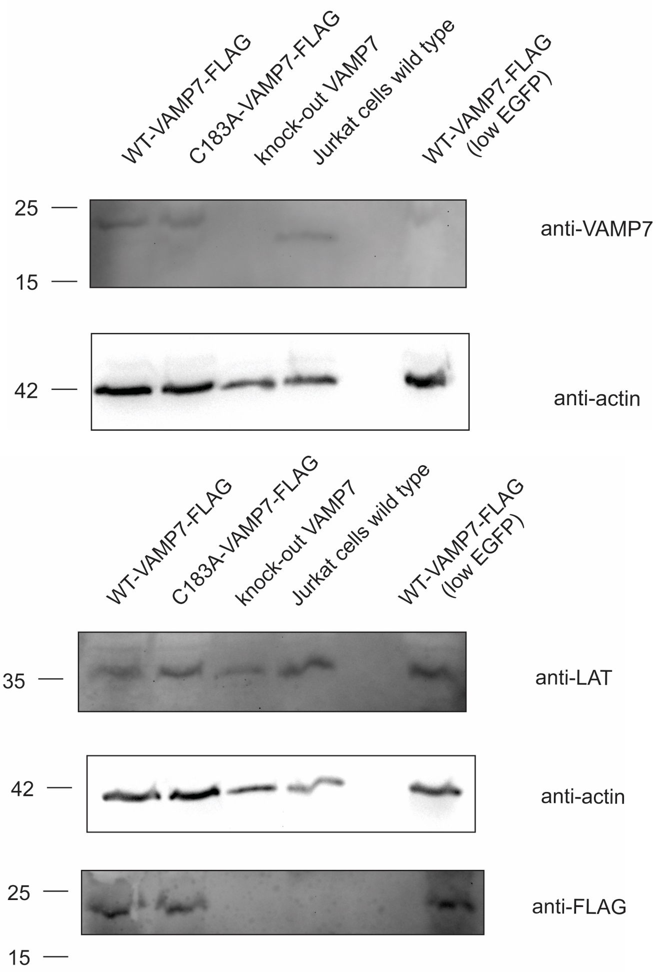


**Supplemental Figure 7.** Confirmation of CRISPR/Cas9-generated VAMP7 knockout and lentivirally transduced VAMP7 WT and C183A construct re-expression Jurkat cell lines. Western blots using anti-VAMP7, anti-FLAG, anti-LAT, and anti-beta-actin antibodies.

Supp. 8

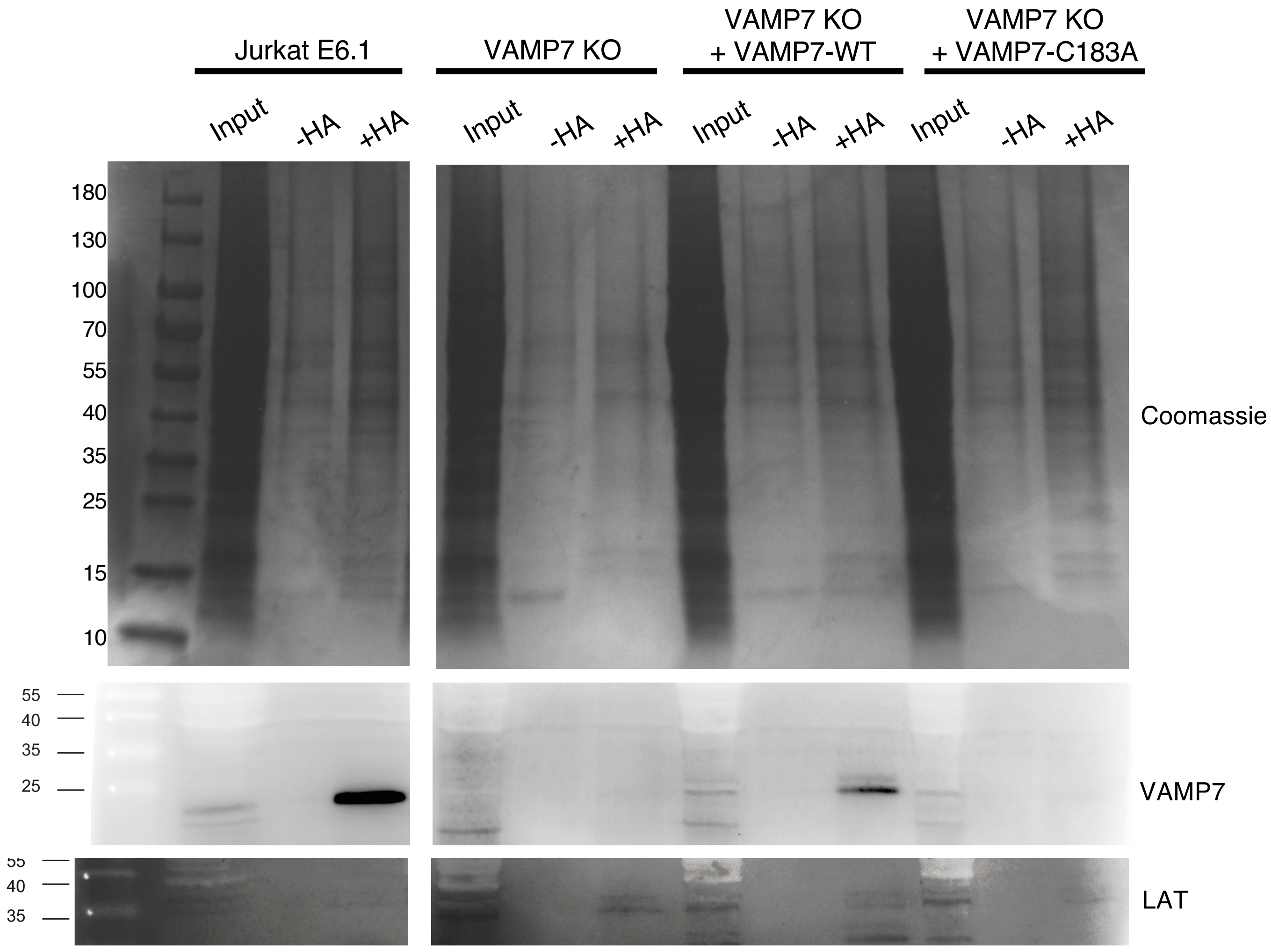


**Supplemental Figure 8.** Confirmation of Cys183 as the site of VAMP7 palmitoylation. Stable VAMP7 KO/re-expression Jurkat cell lines were enriched for palmitoylated proteins using ABE. Western blots using anti-VAMP7 and anti-LAT antibodies confirm that VAMP7-WT is palmitoylated (enriched in +HA), but VAMP7-C183A is not palmitoylated.
